# Theta-gamma transcranial alternating current stimulation enhances motor skill acquisition in healthy young and older adults

**DOI:** 10.1101/2024.10.02.616370

**Authors:** Nishadi N. Gamage, Wei-Yeh Liao, Brodie J. Hand, Philip J. Atherton, Mathew Piasecki, George M. Opie, John G. Semmler

**Author notes:** **Correspondence:** John G Semmler, Ph.D. School of Biomedicine The University of Adelaide Adelaide, South Australia 5005 Australia Telephone: Int.

## Abstract

Theta-gamma transcranial alternating current stimulation (TG tACS) over primary motor cortex (M1) can improve motor skill acquisition in young adults, but the effect on older adults is unknown. This study investigated the effects of TG tACS on motor skill acquisition and M1 excitability in 18 young and 18 older adults. High-definition TG tACS (6 Hz theta, 75 Hz gamma) or sham tACS was applied over right M1 for 20 minutes during a ballistic left-thumb abduction motor training task performed in two experimental sessions. Motor skill acquisition was quantified as changes in movement acceleration during and up to 60 minutes after training. Transcranial magnetic stimulation (TMS) was used to assess changes in M1 excitability with motor-evoked potentials (MEP) and short-interval intracortical inhibition (SICI) before and after training. We found that TG tACS increased motor skill acquisition compared with sham tACS in young and older adults (*P* < 0.001), with greater effects for young adults (*P* = 0.01). The improved motor performance with TG tACS lasted at least 60 minutes after training in both age groups. Motor training was accompanied by greater MEP amplitudes with TG tACS compared to sham tACS in young and older adults (*P* < 0.001), but SICI did not vary between tACS sessions (*P* = 0.40). These findings indicate that TG tACS over M1 improves motor skill acquisition and alters training-induced changes in M1 excitability in healthy young and older adults. TG tACS may therefore be beneficial to alleviate motor deficits in the ageing population.

**Key Points Summary:**

- Theta-gamma transcranial alternating current stimulation (TG tACS) can improve motor function in healthy young adults, but the effect on older adults is unknown.
- We found that TG tACS improved motor skill acquisition with long-lasting effects in healthy young and older adults, but effects were stronger in young adults.
- Transcranial magnetic stimulation showed that TG tACS altered the training-induced changes in motor cortex excitability, but there was no effect of TG tACS on intracortical inhibition in young or older adults.
- Our data suggest that TG tACS represents a promising approach to improve motor skill acquisition throughout the lifespan, and may be beneficial in older patient populations that experience motor or cognitive deficits.

## INTRODUCTION

There is extensive literature on the structural, functional, and biochemical changes in the central nervous system that occur with ageing (see Seidler et al., 2010 for review), but our understanding of how these contribute to the decline in motor skills commonly observed with advancing age is limited. One factor that is thought to contribute to an age-related decline in motor function is a reduced capacity for the brain to change its neural connections, referred to as plasticity (Mahncke et al., 2006, Pascual-Leone et al., 2011, Semmler et al., 2021). In particular, the plasticity of cortical connections plays a major role in the acquisition of new motor skills (Dayan and Cohen, 2011, Sanes and Donoghue, 2000) and altered plasticity has been implicated in reduced motor function in older adults (Ward, 2006, Zapparoli et al., 2022), along with an impaired ability to relearn motor skills following neurological injury (Laaksonen and Ward, 2022). Therefore, it is essential to develop effective interventions to enhance motor system plasticity and improve motor function to promote healthy ageing and recovery from injury.

Non-invasive brain stimulation (NIBS) over primary motor cortex (M1) has been used to manipulate short-term plasticity and improve motor function (Ziemann and Siebner, 2008), but the effects on older adults have been inconsistent (Semmler et al., 2021). For example, when stimulation is applied prior to the motor task (referred to as priming), there is improvement in visuomotor skill learning in older adults with transcutaneous direct current stimulation (tDCS; Fujiyama et al., 2017) and repetitive paired-pulse transcranial magnetic stimulation (TMS; Hand et al., 2023), but not with paired associative stimulation (Opie et al., 2019). An alternative approach has been to perform stimulation to manipulate M1 excitability during the performance of the motor tasks (referred to as gating), with some limited success. For example, anodal tDCS over M1 during motor performance in older adults can improve motor skill acquisition for several different speed-accuracy tasks (Goodwill et al., 2013, Zimerman et al., 2013), and motor adaptation tasks (Panouillères et al., 2015), but not for motor sequence learning (Raw et al., 2016), or visuomotor tracking tasks (Mooney et al., 2019). These inconsistent findings suggest that new NIBS approaches are needed to produce more robust effects on motor skill acquisition in the elderly.

Transcranial alternating current stimulation (tACS) is another NIBS technique that applies small electrical currents to the brain in an oscillatory pattern, which is thought to entrain the membrane potentials of susceptible neurons in the stimulated region at the applied frequency (Krause et al., 2019, Vogeti et al., 2022). Although some effects on motor performance have been observed when tACS is applied over M1 at alpha (10 Hz; Antal et al., 2008) and beta (20 Hz; Pollok et al., 2015) frequencies, there is now strong evidence that tACS in the gamma band (60-90 Hz) modulates M1 plasticity and motor performance. For example, gamma tACS over M1 increases plasticity induced with intermittent theta burst stimulation (iTBS; Guerra et al., 2018), modifies specific aspects of motor behaviour (Joundi et al., 2012, Moisa et al., 2016), and improves different forms of motor skill acquisition (Bologna et al., 2019, Santarnecchi et al., 2017), with these effects thought to be driven by changes in M1 GABA (gamma-aminobutyric acid) mediated intracortical inhibition (Guerra et al., 2018, Nowak et al., 2017). These findings indicate that gamma tACS alters M1 activity during the stimulation period, and these effects are important for increasing plasticity and improving motor behaviour.

Emerging multimodal evidence has shown that high-frequency brain oscillations in the gamma band do not occur in isolation, but are synchronized to low-frequency theta (4-7 Hz) waves during learning (Canolty et al., 2006, Johnson et al., 2017); a process referred to as theta-gamma (TG) phase-amplitude coupling (PAC). Based on these findings, recent studies have applied tACS in a more physiological way by using waveforms that mimic TG coupling and examined the effects on different types of learning. For example, TG tACS applied over the prefrontal cortex has been shown to improve working memory in young (Alekseichuk et al., 2016) and older adults (Reinhart and Nguyen, 2019), and also improves visuomotor learning in older adults (Diedrich et al., 2024). Furthermore, TG tACS has also been applied over M1, where there is an improvement in motor skill acquisition in young adults during a simple ballistic task (Akkad et al., 2021). However, it is not known whether TG tACS applied over M1 is able to improve motor skill acquisition in older adults.

The aim of this study was therefore to examine the effect of TG tACS on motor skill acquisition and M1 excitability in young and older adults. Motor skill acquisition was assessed with a commonly used ballistic thumb abduction task. This task is ideal for examining age-related differences in M1 function and motor skill acquisition as the behavioural improvement is associated with physiological changes in M1 (Muellbacher et al., 2001), the performance of this task is improved with TG tACS in young adults (Akkad et al., 2021), and there are substantial performance decrements for this task in older adults (Rogasch et al., 2009). Neurophysiological changes in M1 were assessed using single and paired-pulse TMS to evaluate alterations in excitatory and inhibitory circuits in the contralateral M1 following TG tACS and motor training. We hypothesised that TG tACS would enhance motor skill acquisition in young and older adults, and this would be accompanied by a greater increase in M1 excitability. In addition, we also hypothesised that effects of TG tACS on motor skill acquisition would be weaker in older adults.

## METHODS

Sample size calculations were performed using simulation-based power estimations on Rstudio (version 2023.12.1 + 402) with ‘*lme4*’ (Bates, 2014) and ‘*simr*’ (Green and MacLeod, 2016) packages. These were based on motor training data obtained from the first five young and older adults (10 participants) and were performed by an unblinded team member that was not involved in data collection. This analysis resulted in a fixed effects coefficient of -0.304 that was used as an unstandardized effect size of TG tACS on motor training in young and older adults, which was lowered by 15% (-0.258) to account for biases in power calculations derived from experimental data (Kumle et al., 2021). Sample size estimations revealed that 24 participants were needed given α < 0.001, β > 0.9 (0.913), which was increased by 50% (36 participants) to ensure robust detection of small–moderate effect sizes.

Eighteen young (8 females; mean age ± SD, 26.3 ± 5.7 years; age range, 18-36 years) and 18 older (12 females; 68.3 ± 5.0 years; age range, 62-78 years) right-handed adults participated in this study. Handedness was assessed by the Edinburgh Handedness Inventory (Oldfield, 1971), and contraindications for TMS were assessed via the TMS safety screening questionnaire (Rossi et al., 2011). Participants were excluded if they had any diagnosed neurological or cardiovascular conditions, a history of concussion or seizure, were pregnant, or were currently on psychoactive medications. Participants who regularly engaged in activities that involve skilled manual dexterity (e.g. musicians) were also excluded. In addition, we assessed participant cognition levels using the Montreal Cognitive Assessment (MoCA) questionnaire (Hobson, 2015), participant physical activity levels using the International Physical Activity Questionnaire- Short form (IPAQ) (Craig et al., 2003), and participant chronotypes using the Morningness-Eveningness questionnaire (MEQr) (Adan and Almirall, 1991). The ethics approval was obtained from the human research ethics committee of the University of Adelaide and conducted according to the *Declaration of Helsinki*. All participants provided written, informed consent prior to the study.

### Experimental design

The current study followed a randomised, double-blind, sham-controlled cross-over design. All participants attended two experimental sessions involving the application of high-definition (HD) theta-gamma tACS (TG tACS) or sham tACS to the right M1 during motor training, with the two sessions separated by at least two weeks to reduce carry-over effects (Bologna et al., 2019). The same experimental procedures were used for both sessions (Figure 1) and were performed by the same experimenter to retain consistency between sessions. For each participant, all experimental sessions were conducted between 11 am and 5 pm at approximately the same time of day to minimise the influence of diurnal variations in circulating cortisol on neuroplasticity (Sale et al., 2008).

**Figure 1.**
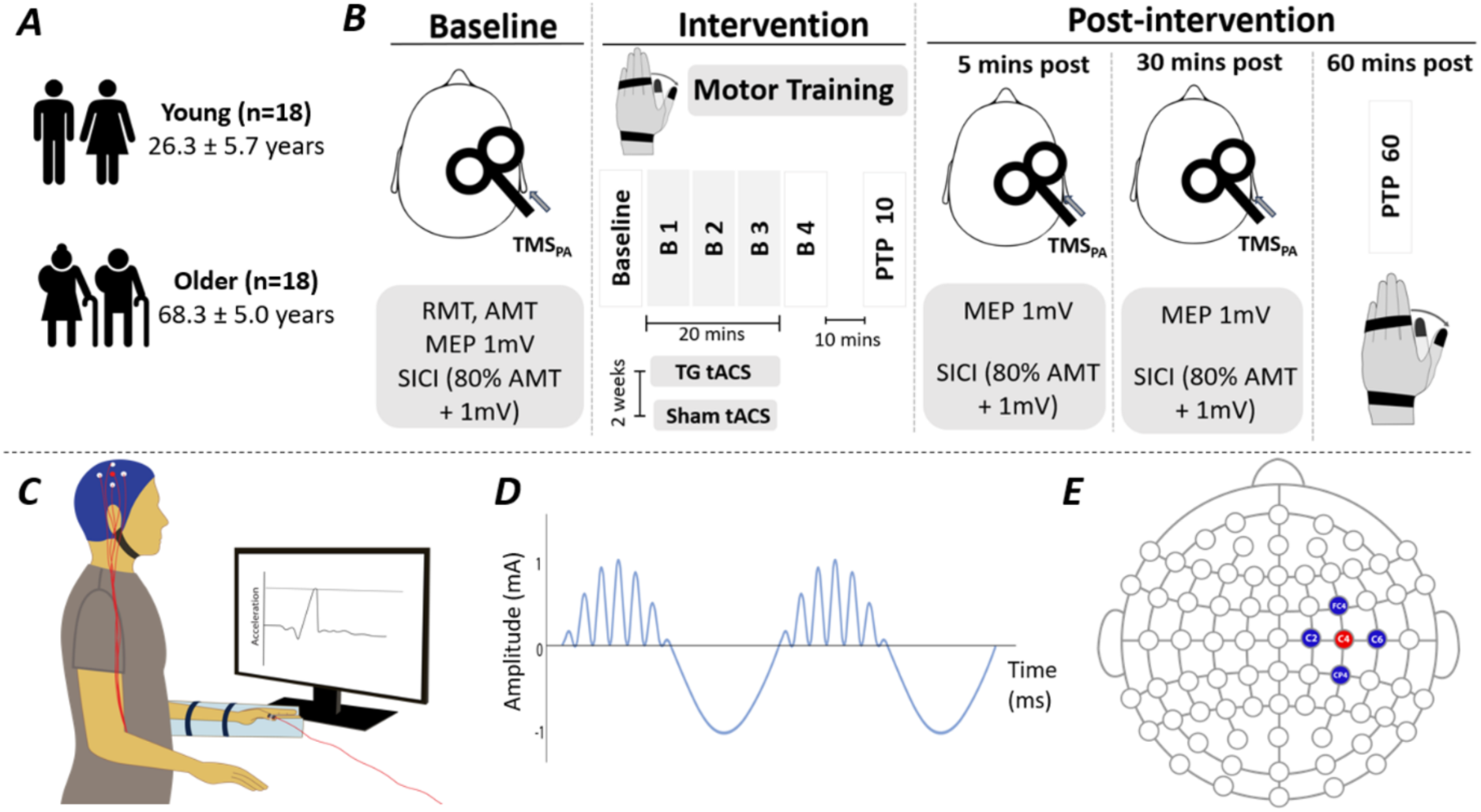
Experimental setup. (A) Study participants. (B) Experimental protocol involving baseline, intervention, and post-intervention measures. RMT, resting motor threshold; AMT, active motor threshold; MEP, motor-evoked potential; SICI, short-interval intracortical inhibition (ISI 2 ms, CS at 80% AMT, and TS at 1mV); B 1-4, motor learning blocks 1-4; PTP, post-training performance. (C) Motor training setup; HD tACS on right M1 while performing ballistic thumb abduction task in the left (non-dominant) hand. (D) Theta-gamma transcranial alternating current stimulation (TG tACS) waveform (sinusoidal 6 Hz theta stimulation at 2 mA peak-to-peak intensity, coupled with bursts of a sinusoidal 75 Hz gamma rhythm amplitude at the positive theta phase). (E) HD tACS electrode placement; anode (red; *C4*) and cathodes (blue; *C2, C6, FC4, CP4*).

### Experimental procedures

In all sessions, participants were seated on a comfortable chair with their non-dominant (left for all) hand relaxed and rested on a table in front of them. Surface electromyography (EMG) was recorded from the abductor pollicis brevis (APB) muscle in the left hand using two Ag-AgCl electrodes placed ∼2 cm apart in a belly-tendon montage on the skin overlying the APB muscle. A third electrode was placed on the radial styloid process of the left hand to ground the electrodes. EMG signals were amplified (300×) and band-pass filtered (20 Hz high pass to 1 kHz low pass) using a CED 1902 signal conditioner (Cambridge Electronic Design, Cambridge, United Kingdom) before being digitised at 2 kHz using a CED 1401 analogue-to-digital converter. Signal noise associated with mains power (50 Hz) was removed using a Humbug mains noise eliminator (Quest Scientific, North Vancouver, Canada). All recorded EMG signals were stored on a computer for offline analysis.

#### Transcranial Magnetic Stimulation (TMS)

The hand area of right M1 was stimulated in all participants using a figure-of-eight branding iron coil (70 mm) connected to two Magstim 200^2^ magnetic stimulators via a BiStim module (Magstim, Dyfed, United Kingdom). The coil was held at an angle of 45° to the midsagittal plane to produce a current flow in a posterior-to-anterior (PA) direction perpendicular to the central sulcus. The stimulation was delivered at a rate of 0.2 Hz with a 10% jitter between trials to avoid anticipation of the stimulus. The identification of APB ‘hotspot’ was determined as the coil position that generated consistent motor-evoked potentials (MEPs) at the lowest stimulation intensity in the contralateral APB muscle. The hotspot was marked using a surgical marker and consistently monitored throughout the experimental session.

Resting motor threshold (RMT) was defined as the smallest stimulus intensity (percentage of maximum stimulation output, % MSO) required to produce a MEP amplitude of ≥ 50 μV in at least 5 out of 10 consecutive trials in the relaxed left APB (Rossini et al., 2015). Following RMT, active motor threshold (AMT) was defined as the lowest stimulus intensity needed to produce a MEP amplitude of ≥ 200 μV in at least 5 out of 10 consecutive trials during a small (∼10% of maximum) contraction of left APB (Rogasch et al., 2009). Then, we determined the single-pulse TMS intensity that produced a MEP amplitude of ∼1.0 mV (range: 0.5 - 1.5 mV) in the resting left APB, when averaged over 20 consecutive trials. The same stimulus intensity was used throughout the experimental session (baseline, 5- and 30-minutes post-training) to assess M1 excitability.

Paired-pulse TMS was used to assess short-interval intracortical inhibition (SICI). This consisted of a sub-threshold conditioning stimulus (CS, 80% AMT) that preceded a suprathreshold test stimulus (TS, 1 mV) by an interstimulus interval (ISI) of 2 ms (Opie and Semmler, 2019). SICI was assessed at all three time points (baseline, 5- and 30-minutes post-training), and each block of SICI consisted of 20 single-pulse and 20 paired-pulse TMS trials delivered in a randomised order, totalling 40 trials. Due to possible changes in M1 excitability following the intervention, we reassessed the baseline single-pulse TMS intensity in a separate TMS block of 20 MEP trials. If the mean MEP amplitude varied from baseline by 30% or greater, the stimulation intensity was readjusted for the SICI block (Nowak et al., 2017), as an intensity producing a larger single-pulse MEP amplitude could induce different levels of intracortical inhibition (Nitsche et al., 2005, Stefan et al., 2002).

#### TG tACS intervention

High-definition (HD) TG tACS was delivered over right M1 via five stimulating HD electrodes (Soterix Medical Inc, New York, United States) arranged in a 4 x 1 HD tACS configuration (4 cathodes and 1 anode) with the anode (active) as the centre electrode (Figure 1E). The electrodes were placed using a stimulating HD cap and connected to an HD-tES stimulator (Soterix Medical MXN-33-HD-tES), which was operated with tACS software on a laptop computer (Soterix Medical, New York, United States). Prior to stimulation, the HD electrodes were attached to the scalp using HD electrolyte gel (Soterix Medical Inc, New York, United States), and the impedance of each electrode was monitored throughout the intervention (< 20 kΩ). Similar to previous work, TG tACS comprised a sinusoidal 6 Hz theta waveform coupled with a 75 Hz gamma burst that was amplitude-modulated by the positive peak of the theta wave. This was applied with a peak-to-peak amplitude of 2 mA (Figure 1D). TG tACS was applied during motor training for 20 minutes (Akkad et al., 2021). During sham tACS, TG tACS (6/75 Hz) was ramped up to 2 mA over 10s and then immediately ramped down again.

Both the experimenter and the participants were blinded to the tACS intervention, which was administered by a second experimenter who was not involved with any other element of data collection or analysis. Following the intervention, participants were asked whether they received real or sham tACS and completed a visual analogue scale (VAS) to assess their perception of pain, intensity, and stimulation localisation.

#### Motor training

In all experimental sessions, participants performed a ballistic thumb abduction task that involved maximizing left thumb acceleration over the training period (Rogasch et al., 2009). The non-dominant (left in all) hand was tested to prevent ceiling effects that might occur in the dominant hand (Akkad et al., 2021). Participants were seated with their left shoulder relaxed and the left elbow flexed to ∼ 90°, while the forearm was maintained in a semi-pronated position with the palm facing inwards. A custom-made plastic splint was used to immobilise the left forearm, wrist, and proximal interphalangeal joints, while the left thumb could move freely. A biaxial accelerometer was connected to the distal phalanx of the thumb to record thumb acceleration in the abduction/adduction (x-axis) and flexion/extension (z-axis) planes (Akkad et al., 2021, Rogasch et al., 2009). Acceleration signals were digitised at 2 kHz with a CED interface system and recorded on a computer for offline analysis.

Participants performed the ballistic thumb abduction training at a rate of 0.4 Hz, paced by a metronome. The training consisted of an initial baseline block (block 0, 30 trials), followed by four training blocks: blocks 1-4 (70 trials each with a 30-second break after trial 35), each separated by a 2-minute rest (no thumb movement) period to avoid fatigue (Akkad et al., 2021). TG tACS or sham tACS was applied during blocks 1-3. Post-training performance was assessed at 10 (PTP 10) and 60 (PTP 60) minutes by performing additional thumb abduction blocks (70 trials each with a 30-second break after trial 35). Participants were continually encouraged to optimise thumb abduction acceleration, while also matching the metronome as accurately as possible (Akkad et al., 2021). To encourage performance improvement, real-time visual feedback of thumb acceleration was presented on a screen in front of the participant. During each break, participants were additionally shown plots of performance over all completed blocks.

#### Maximal compound muscle action potential (M-waves)

The median nerve was stimulated at the wrist to obtain maximal compound muscle action potential (M-waves) in the left APB using electrical stimulation. A constant current stimulator (DS7A, Digitimer, United Kingdom) was used to apply the stimuli, and the cathode of the stimulating bar electrode was positioned distally (Opie and Semmler, 2019). Each stimulus was a square wave pulse of 0.1 ms duration and the stimulator intensity was set at 120% of that required to elicit a maximal response from APB (Opie and Semmler, 2019). The maximal M- wave (*M*max) was recorded as the average of five trials recorded at the beginning of each experimental session. One young participant was unable to tolerate the M-wave assessment.

#### Grooved pegboard task

The grooved pegboard task was used to assess manual dexterity (Reuben et al., 2013). The pegboard contained 25 holes (five-by-five layout), each of which had a ‘key’ shaped profile with a centre circle and radial groove (oriented in a unique direction). Completion of the task requires pegs with matching profiles to be placed in as many holes as possible over a 30-second period. Participants completed three blocks with the left hand, with a 30-second rest between each block. The average number of pegs that were correctly positioned with the left hand was recorded. Twelve young and 15 older participants completed the pegboard task at the start of the first session.

### Data analysis

All EMG data were visually inspected, with any trials having EMG activity exceeding 20 μV peak-to-peak amplitude in the 100 ms prior to TMS excluded from analysis. The MEP and M-wave amplitude were measured peak-to-peak and expressed in mV. SICI was quantified by expressing the MEP amplitude during each paired-pulse trial as a percentage of the mean single-pulse MEP amplitude within the same block. A SICI value of 0% indicates maximum inhibition, whereas 100% represents no inhibition (Opie and Semmler, 2019).

For the ballistic thumb abduction task, acceleration in the abduction/adduction axis was measured in m/s^2^ and was quantified relative to a baseline period 200 ms prior to movement (Rogasch et al., 2009). Trials with a negative value (maximum acceleration < 0 m/s^2^) were removed prior to analysis. In addition, post-training performance (PTP) was assessed by normalizing PTP 10 and PTP 60 acceleration values to the baseline.

### Statistical analysis

Statistical analysis was performed using IBM SPSS (version 28.0) and visualised using Prism (GraphPad software, version 9.0). The distribution of data was visually inspected and assessed using Kolmogorov–Smirnov tests which revealed that the data was non-normally distributed and positively skewed. Therefore, generalised linear mixed models (GLMMs) with Gamma distribution and log link function were used for statistical analyses (Lo and Andrews, 2015). Each model included single-trial data with repeated measures and included random participant intercepts (Barr et al., 2013). The model fit was assessed with the Bayesian Information Criterion (BIC).

#### Baseline measures

Two-factor GLMMs were used to investigate the effects of treatment session (TG tACS, sham tACS) and age group (young, older) on baseline RMT, AMT, TS intensities, single-pulse MEP amplitude, SICI, M-wave, and thumb acceleration.

#### Post training measures

Two three-factor GLMMs were used to investigate the effects of treatment sessions, age groups, and time (baseline, post 5, post 30) on single-pulse MEP amplitude and SICI. A single three-factor GLMM for motor training was used to investigate the effects of treatment sessions, age groups, and training blocks (blocks 0-4) on motor skill acquisition. In addition, a three-factor model was used to investigate the effects of treatment sessions, age groups, and PTP blocks (PTP 10, PTP 60) on post-training thumb abduction. Spearman’s correlation test was performed to assess the association between neurophysiological measures and motor performance. All models assessing MEP, SICI, and thumb acceleration also include participant IPAQ, MoCA, and MEQr scores as covariates to control for variations in participant factors. For models assessing thumb acceleration during motor training, the effect of session order (session 1, session 2) was added to assess whether thumb abduction acceleration improved during the second session. For all models, investigations of main effects and interactions were completed using custom contrasts with Bonferroni corrections, and significance was set at *P* < 0.05. Data for all models are presented as estimated marginal means (EMM) with 95% confidence intervals (95% CI), whereas pairwise comparisons are presented as estimated mean differences (EMD) and 95% CI for an unstandardised measure of effect size.

#### VAS responses

Differences in the perception of TG tACS and sham tACS were investigated by comparing VAS responses (pain, intensity, and localization) using paired sample t-tests, with data presented as mean ± SD.

## RESULTS

All participants completed the experimental sessions with no adverse effects. Age-related differences in participant characteristics are shown in Table 1. Pegboard test scores varied between age groups (*F*1,24 = 8.5, *P* = 0.007), with young adults obtaining higher scores than older adults (both *P* < 0.01). Additionally, chronotype (MEQr values) varied between age groups (*F*1,34 = 5.3, *P* = 0.028), with *post hoc* tests revealing higher MEQr values for older adults compared to young adults (EMD = 1.6, [0.17, 2.94], *P* = 0.029). There were no differences between other characteristics (all *P* > 0.05).

**Table 1.** Participant characteristics in young and older adults.

| Measure | Young | Older | *P* value |
| --- | --- | --- | --- |
| Sample size (F, M) | 18 (8, 10) | 18 (12, 6) | N/A |
| Age (years) | 26.3 [24.3, 28.4] | 68.3 [63.2, 73.9] | < 0.001* |
| Handedness (LQ) | 0.80 [0.72, 0.90] | 0.91 [0.82, 1.01] | 0.109 |
| Pegboard test- L hand | 11.8 [10.3, 13.5] | 9.1 [8.0, 10.3] | 0.007* |
| Physical activity (IPAQ score) | 2461 [1726, 3509] | 2089 [1465, 2979] | 0.512 |
| Cognition (MoCA score) | 28.5 [27.9, 29.1] | 28.8 [28.1, 29.4] | 0.526 |
| Chronotype (MEQr score) | 14.2 [13.3, 15.1] | 15.7 [14.7, 16.8] | 0.028* |

Data presented as EMM [lower 95% CI, upper 95% CI]. LQ, laterality quotient [from -1 (left-handed) to +1 (right-handed)]. IPAQ, International Physical Activity Questionnaire- short form; MoCA, Montreal Cognitive Assessment; MEQr, Morningness-Eveningness questionnaire. Only 26 adults (12 young, 14 older) completed bilateral pegboard tests. \*Significantly different between young and older adults.

Baseline neurophysiological measures are shown in Table 2. Baseline M-wave responses varied between age groups (*F*1,65 = 8.3, *P* = 0.005), and *post hoc* comparisons revealed a greater M-wave response in young adults compared to older adults (EMD = 3.1 mV, [0.87, 5.27], *P* = 0.007). There were no baseline differences for all other neurophysiological measures (all *P* > 0.08). Furthermore, there was no effect of IPAQ, MoCA, and MEQr on MEP amplitude, SICI, and changes in thumb acceleration throughout both experimental sessions (all *P* > 0.05).

**Table 2.**
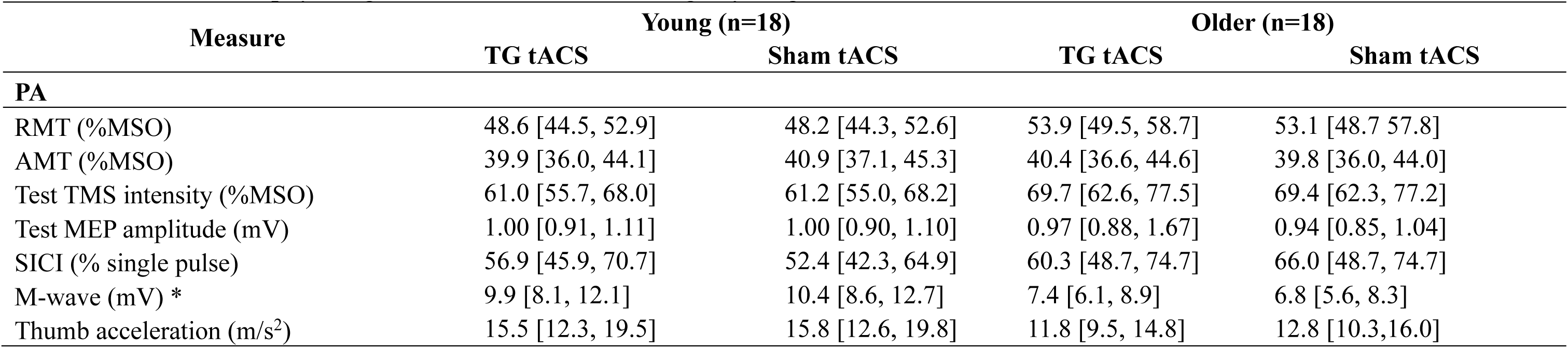
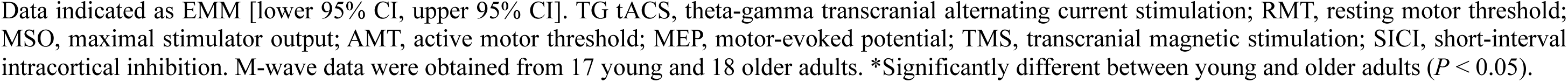
Baseline neurophysiological measures before training in young and older adults.

| Measure | Young (n=18) |  | Older (n=18) |  |
| --- | --- | --- | --- | --- |
|  | TG tACS | Sham tACS | TG tACS | Sham tACS |
| **PA** |  |  |  |  |
| RMT (%MSO) | 48.6 [44.5, 52.9] | 48.2 [44.3, 52.6] | 53.9 [49.5, 58.7] | 53.1 [48.7, 57.8] |
| AMT (%MSO) | 39.9 [36.0, 44.1] | 40.9 [37.1, 45.3] | 40.4 [36.6, 44.6] | 39.8 [36.0, 44.0] |
| Test TMS intensity (%MSO) | 61.0 [55.7, 68.0] | 61.2 [55.0, 68.2] | 69.7 [62.6, 77.5] | 69.4 [62.3, 77.2] |
| Test MEP amplitude (mV) | 1.00 [0.91, 1.11] | 1.00 [0.90, 1.10] | 0.97 [0.88, 1.67] | 0.94 [0.85, 1.04] |
| SICI (% single pulse) | 56.9 [45.9, 70.7] | 52.4 [42.3, 64.9] | 60.3 [48.7, 74.7] | 66.0 [48.7, 74.7] |
| M-wave (mV) * | 9.9 [8.1, 12.1] | 10.4 [8.6, 12.7] | 7.4 [6.1, 8.9] | 6.8 [5.6, 8.3] |
| Thumb acceleration (m/s<sup>2</sup>) | 15.5 [12.3, 19.5] | 15.8 [12.6, 19.8] | 11.8 [9.5, 14.8] | 12.8 [10.3, 16.0] |

Data indicated as EMM [lower 95% CI, upper 95% CI]. TG tACS, theta-gamma transcranial alternating current stimulation; RMT, resting motor threshold; MSO, maximal stimulator output; AMT, active motor threshold; MEP, motor-evoked potential; TMS, transcranial magnetic stimulation; SICI, short-interval intracortical inhibition. M-wave data were obtained from 17 young and 18 older adults. \*Significantly different between young and older adults ( $P < 0.05$ ).

### Thumb acceleration during motor training

Changes in thumb acceleration during motor training (blocks 1-4) are shown in Figure 2. Thumb acceleration during motor training did not vary between age groups (*F*1,19973 = 2.3, *P* = 0.13), but differed between training blocks (*F*3, 19973 = 296, *P* < 0.001), revealing a progressive increase in thumb acceleration from blocks 1 to 4 (all comparisons *P* < 0.001). Changes in thumb acceleration also varied between tACS sessions (*F*1,19973 = 2294, *P* < 0.001), with increased thumb abduction acceleration during the TG tACS session compared to sham tACS (EMD = 59.2% [56.8, 61.7], *P* < 0.001). Furthermore, there was a two-way interaction between age group and training block (*F*3,19973 = 47.8, *P* < 0.001), tACS treatment and training block (*F*3,19973 = 45.7, *P* < 0.001), and tACS treatment and age group (*F*1,19973 = 377, *P* < 0.001; Fig. 2A). In particular, *post-hoc* analysis for the treatment and age group interaction revealed increased thumb acceleration following TG tACS compared to sham tACS for both young (EMD = 83.2% [79.8, 86.7], *P* < 0.001) and older adults (EMD = 35.2% [31.8, 38.7], *P* < 0.001). Moreover, changes in thumb acceleration were increased following TG tACS in young compared to older adults (EMD = 59.2% [13.8, 104.5], *P* = 0.01). There was also a three-factor interaction between tACS treatment, age group, and training block (*F*3,19973 = 4.3, *P* = 0.005; Fig. 2B, 2C). *Post hoc* comparisons showed increased changes in acceleration with TG tACS compared to sham tACS for young and older adults during blocks 1-4 (all *P* < 0.001). Young adults also showed increased changes in acceleration compared to older adults during the TG tACS session for blocks 2-4 (all *P* < 0.02), whereas there were no differences between age groups during sham tACS (all *P* > 0.38). Finally, changes in thumb acceleration differed with session order (*F*1, 19973 = 109, *P* < 0.001), where there were increased changes in acceleration during the first session compared to the second session (EMD = 13.2% [10.76, 15.73], *P* < 0.001).

**Figure 2.**
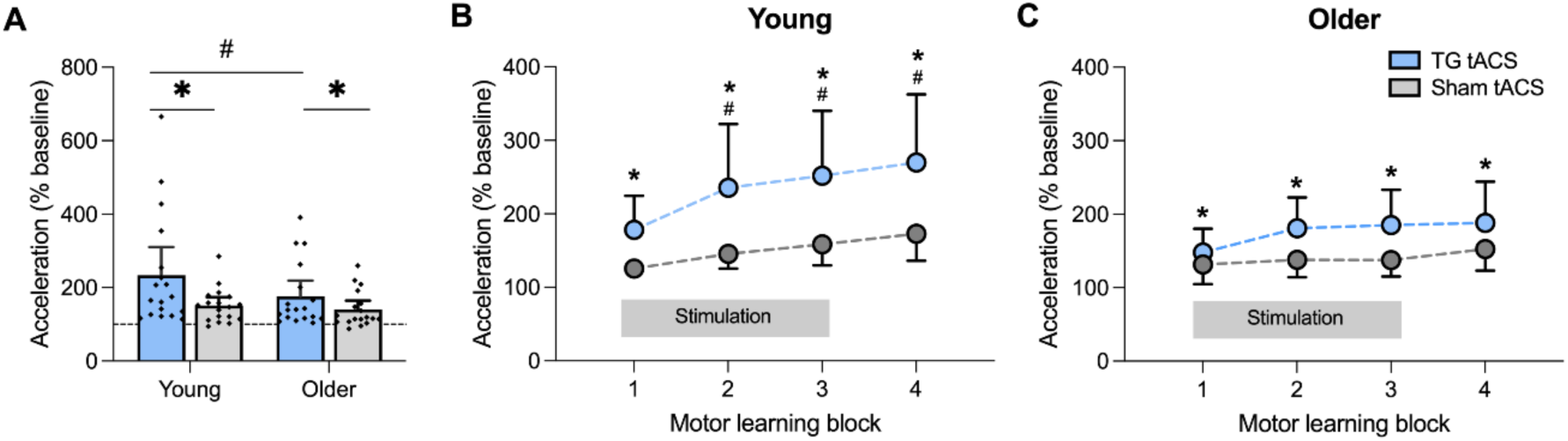
Effect of TG tACS on motor skill acquisition. Data show EMM [± 95% CI] improvement (relative to baseline) in thumb abduction acceleration during motor training (block 1-4). Data show the effect of tACS in different age groups (A), changes in thumb acceleration with training in young adults (B), and changes in thumb acceleration with training in older adults (C). * Significantly different between TG tACS and sham tACS (*P* < 0.001). *^#^* Significantly different compared to older adults (*P* < 0.05).

### Post-training motor skill performance

Changes in thumb acceleration after motor training (PTP) are shown in Figure 3. PTP thumb acceleration differed between tACS sessions (*F*1,10006 = 1076, *P* < 0.001), where thumb abduction acceleration was greater following TG tACS compared to sham tACS (EMD = 53.4% [50.3, 56.6], *P* < 0.001). Thumb acceleration performance also varied between age groups (*F*1,10006 = 5.1, *P* = 0.024), with increased changes in acceleration in young compared to older adults (EMD = 51.5% [6.7, 96.4], *P* = 0.024). Additionally, changes in PTP thumb acceleration differed between PTP blocks (*F*1,10006 = 162, *P* < 0.001), with reduced changes in thumb acceleration during PTP 60 compared to PTP 10 (EMD = 20.7% [17.5, 23.9], *P* < 0.001). There was an interaction between age group and PTP block (*F*1,10006 = 11.4, *P* < 0.001) and tACS treatment and PTP blocks (*F*1,10006 = 40.3, *P* < 0.001; Fig, 3A), which showed increased changes in PTP thumb acceleration following TG tACS compared to sham tACS at PTP 10 (EMD = 63.8% [59.3, 68.3], *P* < 0.001) and PTP 60 (EMD = 43.1% [38.6, 47.6], *P* < 0.001).

**Figure 3.**
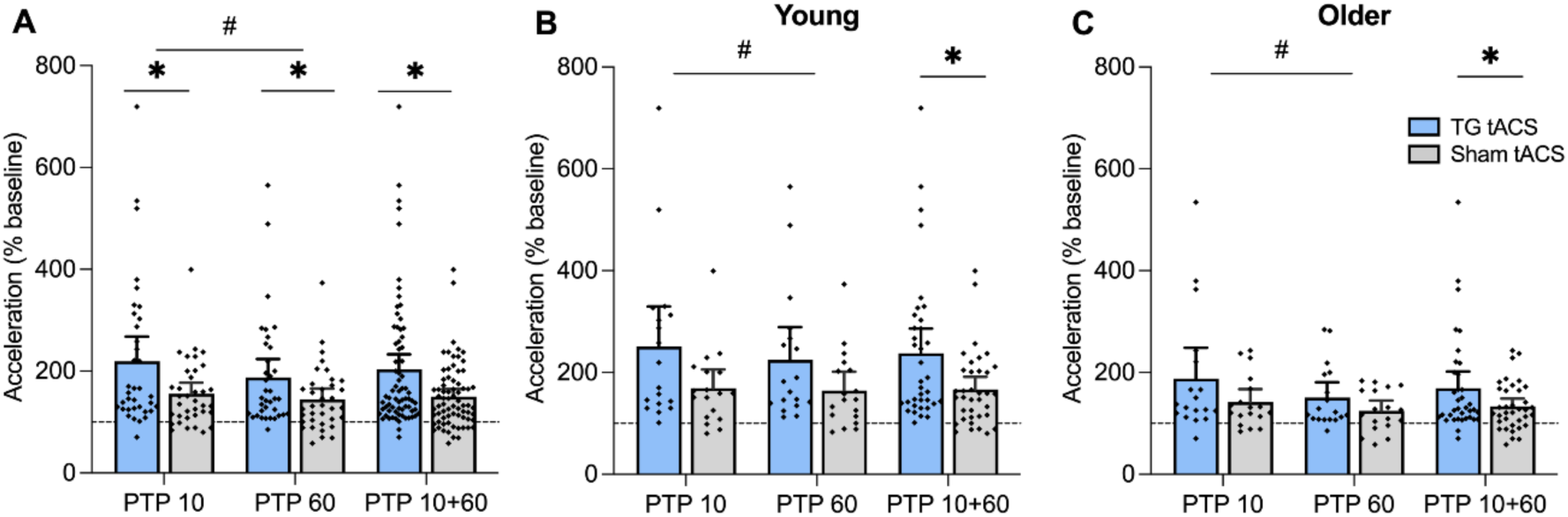
Effect of motor training and TG tACS on thumb acceleration at post-training performance (PTP). Data show EMM [± 95% CI] thumb acceleration (relative to baseline) in all subjects (A), young adults (B), and older adults (C) obtained 10 minutes (PTP 10) and 60 minutes (PTP 60) after training, and with both time points combined (PTP 10+60). *^#^* Significant difference between PTP 10 compared to PTP 60 (*P* < 0.001). * Significant difference between TG tACS and sham tACS (*P* < 0.001).

Additionally, changes in PTP thumb acceleration at PTP 10 was greater compared to PTP 60 for TG tACS (EMD = 31.1% [26.6, 35.6], *P* < 0.001) and sham tACS (EMD = 10.4% [5.9,], *P* < 0.001). Furthermore, there was an interaction between age group and PTP block (*F*1,10006 = 11.4, *P* < 0.001; Fig. 3B, 3C), where young adults showed increased changes in PTP acceleration at PTP 10 (EMD = 46.0% [1.1, 91.0], *P* = 0.045) and PTP 60 (EMD = 57.0% [12.1, 102.0], *P* = 0.013) compared to older adults. Moreover, changes in PTP thumb acceleration were increased during PTP 10 compared to PTP 60 for both young (EMD = 15.2% [10.7, 19.7], *P* < 0.001) and older (EMD = 26.2% [21.7, 30.8], *P* < 0.001) adults. There were no significant three-way interactions (*P* = 0.81).

### Effect of motor training and TG tACS on MEP Amplitude

MEP amplitude obtained before and after training (at the same TMS intensity) for each tACS session are shown in Figure 4. MEP amplitude did not differ between age groups (age effect, *F*1,3996 = 1.7, *P* = 0.187), but varied between tACS treatments (treatment effect, *F*1,3996 = 36.8, *P* < 0.001), where TG tACS resulted in increased MEP amplitude compared to sham tACS (EMD = 0.13 mV [0.08, 0.17], *P* < 0.001). In addition, MEP amplitude differed between the timepoints (time effect, *F*2,3996 = 8.1, *P* < 0.001), with increased MEP amplitude 5 minutes following the intervention compared to 30 minutes (EMD = 0.06 mV [0.01, 0.12], *P* = 0.02). There was a significant interaction between tACS treatment and timepoint (*F*2,3996 = 7.0, *P* < 0.001; Fig. 2A), with *post hoc* analysis showing increased MEP amplitude following TG tACS compared with sham tACS at post-5 minutes (EMD = 0.16 mV [0.09, 0.23], *P* < 0.001) and post-30 minutes (EMD = 0.19 mV [0.12, 0.25], *P* < 0.001). MEP amplitude following sham tACS was also decreased at post-5 minutes (EMD = 0.10 mV [0.03, 0.18], *P* = 0.006) and post- 30 minutes (EMD = 0.18 mV [0.10, 0.26], *P* < 0.001) compared to baseline. There was also an interaction between tACS treatment and age groups (*F*1,3996 = 7.5, *P* = 0.006), which revealed increased MEP amplitude following TG tACS compared to sham in young (EMD = 0.07 mV [0.01, 0.13], *P* = 0.02) and older adults (EMD = 0.17 mV [0.11, 0.23], *P* < 0.001). There were no other interactions (all *P* > 0.114).

**Figure 4.**
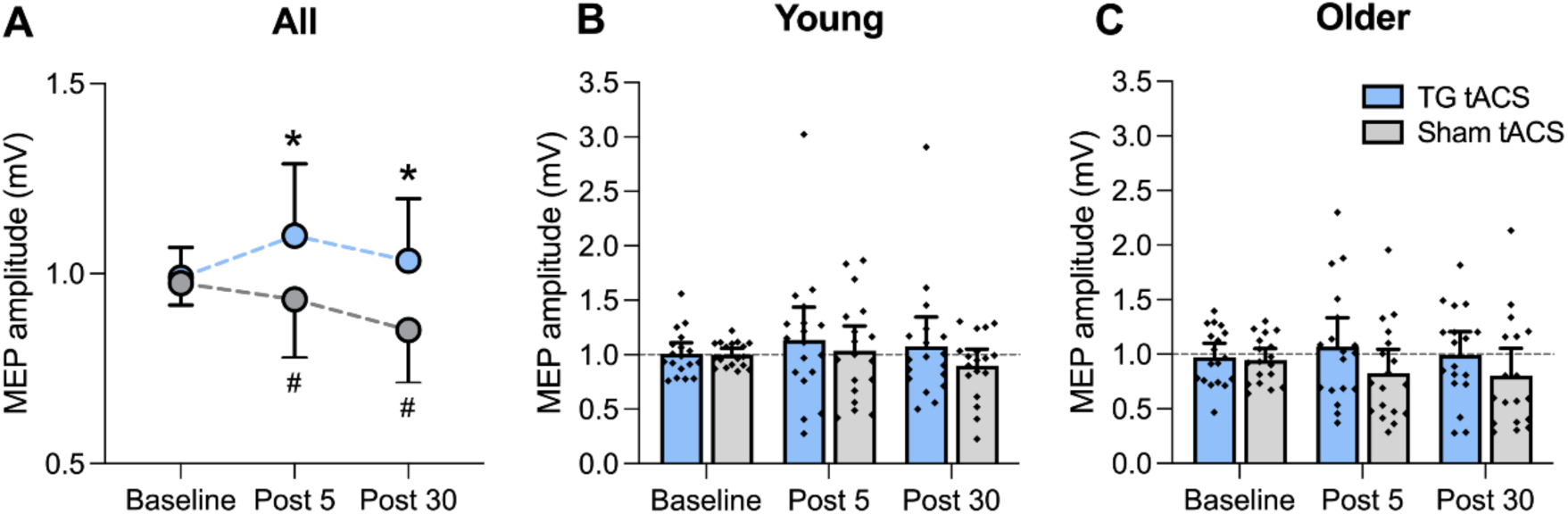
Effect of motor training and TG tACS on MEP amplitude. Data show EMM [± 95% CI] test MEP amplitude in all participants (A), young adults (B), and older adults (C). * Significantly different compared to sham tACS (*P* < 0.001). *^#^* Significantly different compared to baseline (*P* < 0.01).

### Effect of motor training and TG tACS on SICI

Pre- and post-training SICI for each tACS session are shown in Figure 5. SICI did not vary between tACS treatments (*F*1,4061 = 0.70, *P* = 0.40) or age groups (*F*1,4061 = 0.78, *P* = 0.38), but differed between timepoints (*F*2,4061 = 7.5, *P* < 0.001), with *post hoc* comparisons revealing decreased SICI at post-5 minutes (EMD = 8.0% [2.5, 12.9], *P* = 0.001) and post-30 minutes (EMD = 5.8% [1.0, 10.5], *P* = 0.01) relative to baseline (Figure 5A). Moreover, there was a significant interaction between age group and tACS treatment (*F*1,4061 = 16.2, *P* < 0.001), with *post hoc* tests showing lower SICI for young during the TG tACS session compared to sham (EMD = 5.1% [0.56, 9.72], *P* = 0.03), whereas SICI was greater for older adults during the TG tACS session compared to sham (EMD = 8.9% [3.5, 14.3], *P* = 0.001). There were no other interactions (all *P* > 0.09). The mean single pulse MEP amplitudes from which SICI was obtained did not differ between tACS treatments (*P* = 0.65), but were influenced by age group (*F*1,4067 = 4.4, *P* = 0.04) and time (*F*2,4067 = 9.6, *P* < 0.001). *Post hoc* analyses showed that single-pulse MEP amplitudes (within the SICI block) were greater for young compared with older adults (EMD = 0.13 mV [0.01, 0.26]), and greater post-5 minutes (EMD = 0.06 mV [0.01, 0.12], *P* = 0.02) and post-30 minutes (EMD = 0.11 mV [0.05, 0.17], *P* = 0.001) relative to baseline.

**Figure 5.**
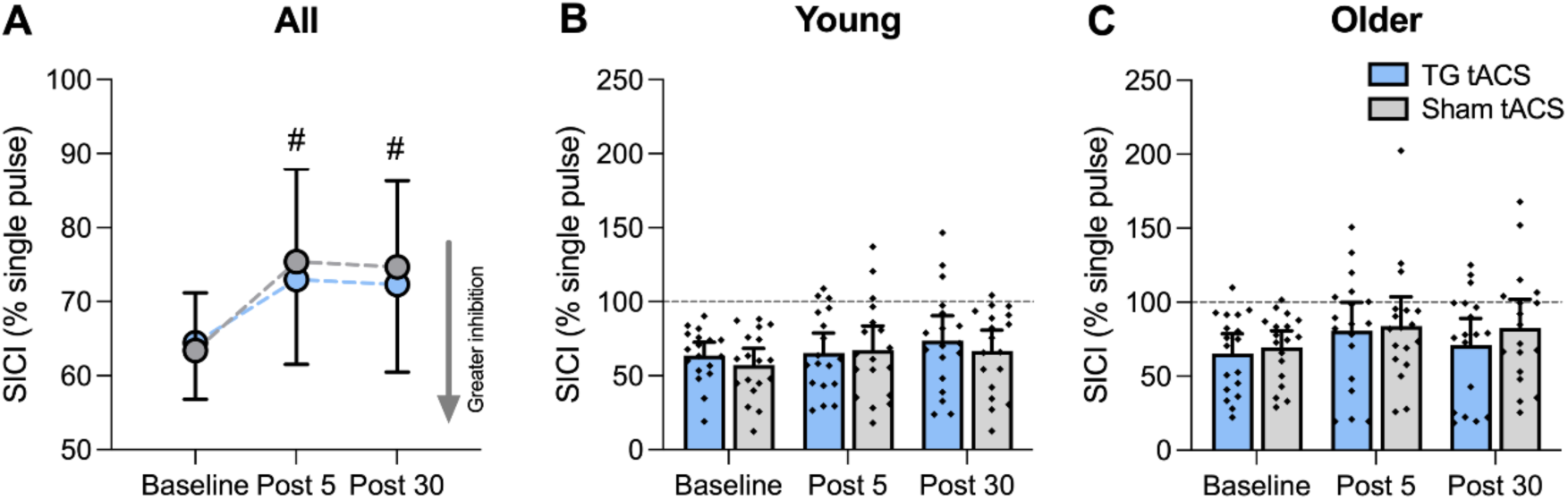
Effect of motor training and TG tACS on SICI. Data show EMM [± 95% CI] SICI (% single pulse) in all participants (A), young adults (B), and older adults (C). *^#^* Significantly different compared to baseline for TG tACS and sham tACS combined (*P* < 0.05).

### Pain, intensity, localization (VAS score), and HD tACS blinding of participants

Responses to VAS and blinding questionnaires are presented in Table 3. There were no differences in VAS scores for the level of pain, intensity, and HD tACS stimulation localization between tACS sessions (all *P* > 0.10). During the TG tACS session, 53% (n = 18) of participants stated the intervention correctly, while 32% (n = 11) stated it incorrectly. In contrast, during the sham tACS session, 24% (n = 8) stated the intervention correctly, 61% (n = 20) of the participants stated it incorrectly, and 15% (n = 5) of participants were unable to differentiate between the two stimulation types in both sessions.

**Table 3.**
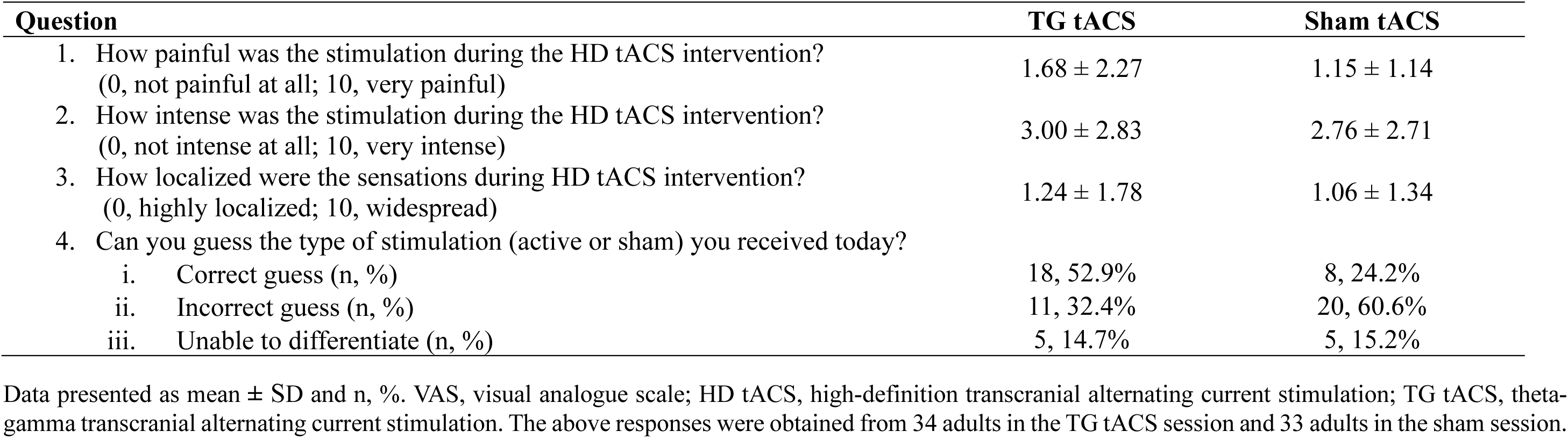
Comparison of VAS responses (mean ± SD) and HD tACS intervention blinding of participants (n, %) between sessions. Question TG tACS Sham tACS.

| Table 3. Comparison of VAS responses (mean ± SD) and HD tACS intervention blinding of participants (n, %) between sessions. |  |  |
| --- | --- | --- |
| Question | TG tACS | Sham tACS |
| 1. How painful was the stimulation during the HD tACS intervention?<br>(0, not painful at all; 10, very painful) | 1.68 ± 2.27 | 1.15 ± 1.14 |
| 2. How intense was the stimulation during the HD tACS intervention?<br>(0, not intense at all; 10, very intense) | 3.00 ± 2.83 | 2.76 ± 2.71 |
| 3. How localized were the sensations during HD tACS intervention?<br>(0, highly localized; 10, widespread) | 1.24 ± 1.78 | 1.06 ± 1.34 |
| 4. Can you guess the type of stimulation (active or sham) you received today? |  |  |
| i. Correct guess (n, %) | 18, 52.9% | 8, 24.2% |
| ii. Incorrect guess (n, %) | 11, 32.4% | 20, 60.6% |
| iii. Unable to differentiate (n, %) | 5, 14.7% | 5, 15.2% |

Data presented as mean ± SD and n, %. VAS, visual analogue scale; HD tACS, high-definition transcranial alternating current stimulation; TG tACS, theta-gamma transcranial alternating current stimulation. The above responses were obtained from 34 adults in the TG tACS session and 33 adults in the sham session.

### Correlation between neurophysiological measures and skill acquisition

There were no significant correlations between neurophysiological measures (MEP amplitudes and SICI) and thumb acceleration performance in young or older adults for both tACS sessions (all *P* > 0.05).

## DISCUSSION

It has previously been shown that TG tACS applied over M1 can improve motor skill acquisition of a ballistic thumb abduction task in young adults (Akkad et al., 2021). In this previous study, TG tACS was shown to be most effective when the gamma oscillation (75 Hz) was applied at the peak of the theta wave (6 Hz), and this finding was replicated in an independent sample from the same study showing that this effect lasted at least one hour (Akkad et al., 2021). Using the same motor task and training protocol but with more focal HD tACS (compared with standard tACS) over M1, we show a substantial improvement in motor skill acquisition in young participants with TG tACS (increase of 134% *vs.* 51% for sham tACS) that was also long-lasting (> 60 mins), indicating that this is a robust effect. In addition, we show for the first time that TG tACS can improve motor skill acquisition in healthy older adults. During training for the ballistic task in older adults, thumb abduction acceleration increased by 76% after TG tACS compared with 40% after sham tACS, showing an almost twofold improvement in ballistic thumb abduction acceleration with TG tACS in older adults. TG tACS was also accompanied by a long-lasting improvement of task performance at 10 (increase of 88% *vs.* 42% for sham tACS) and 60 mins (increase of 51% *vs.* 25% for sham tACS) after motor training in older adults.

In contrast to using separate groups in the previous study (Akkad et al., 2021), participants in our protocol performed both sessions in random order that were separated by at least two weeks. While we found similar baseline performance for each session, covariate analyses identified a significant order effect, where there was greater improvement in thumb abduction acceleration in the first compared with the second session. This was unexpected, as it has previously been reported that there was no session order effect when this task was performed at least one week apart (Bologna et al., 2019). Nonetheless, our random allocation suggests that session order did not contribute to the greater effects observed with TG tACS. Furthermore, participants were unable to differentiate between the two tACS conditions, with similar VAS scores for the level of pain, intensity, and stimulation localization, suggesting successful participant blinding to the stimulation condition.

The effects of TG tACS on motor skill acquisition were reduced in older adults, and several factors could contribute to this difference. First, we used the same tACS parameters (intensity, frequency) in young and older adults, but the optimal parameters may be altered with advancing age. We did not obtain structural magnetic resonance images for our participants, so we were unable to estimate the effective current reaching the cortex, which could be reduced in older adults (when stimulating at the same intensity) due to cortical atrophy (Antal et al., 2017, Freitas et al., 2011). However, it has previously been shown with gamma tACS that the current density reaching M1 was similar in young and older adults when using the same tACS intensity (Guerra et al., 2021). Furthermore, brain oscillations in sensorimotor areas are altered with advancing age (Ishii et al., 2018), including age-related changes in TG coupling (Reinhart and Nguyen, 2019, Zhang et al., 2021), so it is possible that the effectiveness of tACS at the frequencies used here (6 Hz theta, 75 Hz gamma) may not be optimal in older adults. Second, it is commonly accepted that M1 plasticity declines with advancing age (Freitas et al., 2013, Semmler et al., 2021, Zimerman and Hummel, 2010), with this capacity thought to be a key contributor to the early stages of motor skill learning (Classen et al., 1998, Dayan and Cohen, 2011). This reduced plasticity could occur through morphological, neurochemical, and neurophysiological changes in the M1 network in older adults (Seidler et al., 2010), resulting in reduced motor skill learning with age (Voelcker-Rehage, 2008). Finally, it is possible that the maximum capacity for thumb acceleration is reduced in older adults (i.e. lower ceiling effect) due to age-related changes in neuromuscular function, which limits the potential improvement with TG tACS. For example, the ability to perform ballistic tasks could be reduced in the elderly because of a decline in the proportion of muscle occupied by type II fibers (Klein et al., 2003), which are likely to make the biggest contribution to thumb abduction acceleration, particularly at the end of training when acceleration is greatest. Furthermore, higher motor unit firing rates during rapid contractions contribute to an increase in contraction speed (Van Cutsem et al., 1998), and maximum firing rates are reduced in older adults during rapid contractions (Klass et al., 2008), which could limit the performance of ballistic tasks in the elderly.

Gamma oscillations that are phase synchronised to the peak of theta waves (i.e. TG phase-amplitude coupling; TG PAC) are known to be an important mechanism for neuronal processing in the hippocampus (Colgin, 2015), where it has been linked with working memory in humans (Alekseichuk et al., 2016, Daume et al., 2024). These observations also extend to the motor system (Canolty et al., 2006), where TG PAC over motor brain regions has been positively associated with motor learning success (Dürschmid et al., 2014), and in motor recovery following brain-computer interface therapy in chronic stroke patients (Rustamov et al., 2022). As demonstrated previously, TG tACS over M1 can improve simple motor skill acquisition in young adults (Akkad et al., 2021), and we now show that TG tACS is effective at modulating simple motor skill acquisition in healthy older adults. These findings suggest that specific features of M1 oscillations are likely to play a role in ballistic motor skill acquisition throughout the healthy lifespan. However, these findings in the motor system are specific to the simple ballistic task used here. This is a widely used experimental task where the explicit goal is to maximise thumb acceleration in the abduction direction, and it results in substantial improvements with practice that are largely dependent on physiological changes in M1 (Muellbacher et al., 2001). Nonetheless, it is currently unknown if TG tACS can modulate motor skill acquisition during more complex visuomotor tasks that more closely represent real-world motor learning requirements (e.g. reaching and grasping), which usually involve coordinated neuronal activity across a distributed network of brain regions involving attentional, memory, visual and motor systems (Dayan and Cohen, 2011). Furthermore, these findings are specific to healthy older adults who demonstrated only minor declines in baseline motor function (Pegboard scores), were moderately active (IPAQ score), and had normal cognitive function (MoCA scores), so the effects of TG tACS may be different in frail older adults, or older patients with neurological injury. This is supported by recent evidence showing that TG tACS over M1 during a more complex motor task impaired motor skill acquisition in chronic stroke survivors (Grigutsch et al., 2024). Taken together, these findings suggest that TG tACS enhances ballistic motor skill acquisition in healthy young and older adults, and future studies should explore the effects of TG tACS on different motor tasks and neurological conditions that have an increased incidence with advancing age.

Motor skill acquisition with the ballistic thumb abduction task is usually accompanied by a post-training increase in MEP amplitude (Muellbacher et al., 2001, Ziemann et al., 2001), which is thought to represent training-dependent plasticity involving LTP-like mechanisms in cortical circuits (Sanes and Donoghue, 2000), although spinal circuits may also be involved in this task (Giesebrecht et al., 2012). In contrast, our data show a reduction in MEP amplitude after motor training (with sham tACS) in young and older adults. This unexpected finding could be due to the extended duration of training undertaken in our study. An increase in MEP amplitude has previously been shown when training is undertaken with this task for shorter durations (Rosenkranz et al., 2007), and we performed a large number of movement trials that lasted ∼25 mins, which could result in a reversal of the expected MEP response due to homeostatic metaplasticity (Hassanzahraee et al., 2020). Furthermore, this effect could be due to the training structure (Lage et al., 2015), where multiple training blocks were performed that were interspersed with rest periods (i.e. interleaved training), as opposed to previous studies showing increased MEPs with less blocks and more repetitive training (Cirillo et al., 2010, Rogasch et al., 2009). Nonetheless, the reduced M1 excitability after motor training was altered with TG tACS, either by interfering with the neurophysiological processes that generated the MEP depression, or by augmenting training-related MEP facilitation through plasticity-related mechanisms. One way this can occur is through the online entrainment of membrane potentials (with TG tACS) that interact with the timing of pre- and post-synaptic potentials during voluntary motor activity (Vogeti et al., 2022), resulting in increased neural synchronisation, which is important for inducing synaptic plasticity (Uhlhaas et al., 2010, Womelsdorf et al., 2007).

A reduction in GABAergic inhibition can also play an important role in training-dependent plasticity and motor skill acquisition (Ziemann et al., 2001), and this can be quantified with paired-pulse TMS to assess SICI (Ziemann et al., 1996). Several studies have shown a reduction in SICI after ballistic thumb movements (Liepert et al., 1998, Rosenkranz et al., 2007), reflecting GABAergic disinhibition in M1 following motor skill acquisition. We found reduced SICI after motor training in young and older adults, which is consistent with these previous findings. However, the magnitude of this change was not different between stimulus conditions, suggesting that TG tACS did not influence training-related GABAergic inhibition. Previous studies have shown that gamma tACS over M1 decreases SICI when assessed during stimulation (Guerra et al., 2018, Nowak et al., 2017), with this online change shown to be predictive of performance on a subsequent motor learning task (Nowak et al., 2017). We were unable to quantify SICI during TG tACS due to the concurrent performance of the motor task, so it is currently unknown whether TG tACS (as opposed to gamma tACS alone) modulates SICI. Furthermore, we could not provide direct evidence of entrainment of brain oscillations with TG tACS, as electroencephalographic recordings are confounded by large stimulation artefacts (Vogeti et al., 2022). Therefore, the neurophysiological mechanisms that contribute to improved motor skill acquisition with TG tACS in healthy young and older adults remain to be determined.

In conclusion, we have shown that TG tACS can enhance motor skill acquisition in healthy young and older adults. These improvements in performance were long-lasting (> 60 mins) and stronger in young compared with older adults. In addition, TG tACS altered training-induced changes in M1 excitability, but did not influence changes in GABAergic inhibition in both age groups. These findings suggest that TG tACS represents a promising approach to improve motor function throughout the lifespan in healthy individuals. To better understand the rehabilitation potential of TG tACS, future research will need to explore the efficacy of different TG tACS protocols with more complex motor tasks and in neurological conditions that have an increased incidence in the ageing population.

## Data Availability Statement

Data from this study will be made available upon reasonable request to the corresponding author.

